# Minding the gap: collective determinants of multiscale structure across interacting bacterial colonies

**DOI:** 10.64898/2026.04.30.721914

**Authors:** Jacob Moran, Madeline Shay, Michael Hinczewski, Suraj Shankar, Kevin B. Wood, Robert J. Woods, Luis Zaman

**Affiliations:** Department of Ecology and Evolutionary Biology, University of Michigan; Department of Cellular and Molecular Biology, University of Michigan; Department of Physics, Case Western Reserve University; Department of Physics, University of Michigan; Department of Biophysics, University of Michigan; Department of Internal Medicine, University of Michigan; Department of Ecology and Evolutionary Biology, Center for the Study of Complex Systems, University of Michigan

## Abstract

A bacterial colony rarely exists in isolation – in natural habitats, colonies interact to form spatially structured communities across length and time scales. Eco-evolutionary feedbacks link these scales, such that structures at one level can influence another. A commonly observed multi-colony trait is the formation of a stable gap between growing colonies (even when isogenic) but, despite its ubiquity, the underlying mechanisms are poorly understood. To address this, we develop a high-throughput robotic platform to track population dynamics across spatially extended networks of colonies. We show that isogenic colonies of the opportunistic pathogen *Enterococcus faecalis* spontaneously form stable gaps when inoculated beyond a critical distance apart. By combining quantitative modeling and experiments, we show that both nutrient competition and direct growth inhibition control colony morphology and expansion of interacting colonies. We identify distinct regimes of gap formation, relating intra- and inter-colony spatial patterns to ecological interactions mediated at the cellular scale. Together, our results suggest that antagonism, even between isogenic populations through self-inhibition, is likely a common behavior of bacterial species in general.

---

Microbes colonize surfaces by forming biofilms or other aggregates with complex spatiotemporal organization spanning multiple scales. These structures emerge from collective processes that bridge cell and colony scale interactions [1–5], with important functional consequences at the ecosystem level, such as in agriculture and the global climate [6]. While substantial progress has been made in understanding the formation of individual biofilms [7– 16], most microbial communities do not exist in isolation: even physically separated colonies can interact at a distance through long-ranged cues (e.g., electrical, metabolic, and chemical signals [17–20]). But how such interactions within and between populations control microbial growth, form, and fitness remains poorly understood.

One such widely observed multiscale behavior, even between clonal populations (“sibling rivalry”), is the conditional merging of spatially separated colonies: colonies initiated at sufficient proximity eventually fuse into a cohesive unit, but those seeded farther apart remain segregated leaving a persistent cell-free gap in between (Fig. 1A). This phenomenon has been reported across strains and species, for motile and immotile cells [21– 28, with similar patterns observed *in-vivo* as well, e.g., in the zebrafish gut [29]. Simple arguments based on nutrient competition and geometric constraints [30, 31], or contact-based recognition [32, 33] have been proposed to explain spatial partitioning of interacting colonies, but are insufficient to capture gap formation, especially in isogenic systems. Recent studies have identified diffuse inhibitory interactions [23, 34, 35] as a mechanism for gap formation in some motile bacteria, highlighting the importance of antagonistic interactions between colonies [21, 28, 36–38] in structuring spatial patterns with fitness consequences. But what are the minimal ingredients and general principles that give rise to this multi-colony structure?

**FIG. 1.**
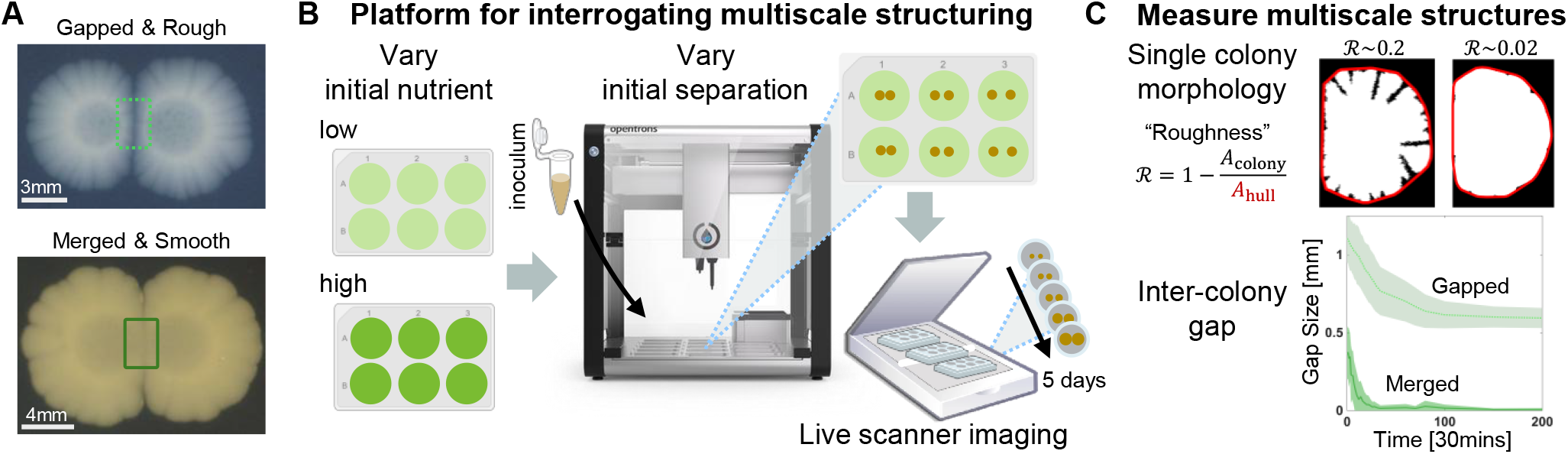
Automated experimental platform for interrogating spatial structure formation in interacting microbial communities. **A:** Pairs of *E. faecalis* colonies can exhibit a range of inter-colony interfaces and intra-colony morphologies. Depending on initial conditions, interacting colonies can individually be either smooth or rough, and can either merge or form a gapped region with no growth (bottom image versus top image, respectively). Images are of colonies after 5 days of expansion grown on agar supplied with either 0.25x (top image) or 1x (bottom image) of standard nutrient concentrations (see Methods). **B:** Pipeline developed to study multi-colony interactions and identify key patterning factors, such as nutrient amount and colony arrangement. Agar plates are prepared with varying amounts of initial nutrient and inoculated with pairs of colonies using an Opentrons™ OT-2 pipetting robot to spot at varying initial distances. Colonies are grown at 37^*°*^C for 5 days and imaged using programmed flatbed document scanners (see Methods; image from Opentrons Labworks, Inc.). **C:** We choose two simple traits (at the final time-point) to quantify single- and multi-colony morphology: (1) colony roughness ℛ measuring the area fraction of the convex hull (outlined in red) that is not filled by the segmented colony pixels (white; see text), and (2) the inter-colony separation (gap size) that measures the average edge-to-edge distance between segmented colonies at closest approach (see Methods for details; errorbars represent standard deviation in gap size within averaging window). Across our experimental runs, we find that colony-pairs either merge within the first 2 days or reach a stable gap size within 3 to 4 days. The examples of roughness and gap trajectories shown here are measured from the examples presented in (**A**).

Here we report the observation of sibling rivalry and gap formation in *Enterococcus faecalis* (Fig. 1A), an immotile, opportunistic pathogen that naturally inhabits the human gut. To map out the multiscale organization of interacting colonies in space, we develop a highthroughput experimental pipeline that combines opensource robotics with automated scanner imaging to track colony networks over time (Fig. 1B,C). By integrating quantitative experiments with a theoretical model, we show that intra- and inter-colony structure organizes into four distinct qualitative phases defined by colony morphology and gap formation. Our model identifies both resource competition and growth inhibition as sufficient mechanisms for gap formation, but their distinct quantitative phenotypes reveal that direct growth inhibition is necessary to capture the full range of experimental observations. We experimentally test this prediction using a spent-agar assay and show that *E. faecalis* colonies produce inhibitory factors that suppress subsequent growth.

Altogether, our results establish a quantitative framework for linking cell-scale interactions to emergent structure across scales, and provide simple diagnostics for identifying dominant interactions from spatial data. Our findings suggest that antagonistic interactions, including self-inhibition, may be a widespread and functionally important feature of microbial communities.

## I. RESULTS

### A. Platform for surveying cross-scale organization in multi-colony networks

Pairs of isogenic colonies of *E. faecalis* exhibit sibling rivalry, i.e., colonies merge when initially seeded sufficiently close together, but form a stable gap with no growth when initialized far apart (Fig. 1A, bottom vs. top). This inter-colony behavior is also accompanied by a range of colony front morphologies, from smooth to rough, depending in part on nutrient availability. To quantitatively probe how these structures form for varied experimental conditions (e.g., initial separation, nutrient conditions, inoculum characteristics) requires a high-throughput platform with precision control and measurements.

We tackle this challenge by developing an automated experimental and imaging pipeline shown in Fig. 1B. We prepare agar plates with varying initial nutrient levels (concentration of brain heart infusion media, [BHI]) and spot inoculum cultures onto the plates using an Opentrons OT-2 liquid handling robot, enabling precise control over initial inter-colony separations. To monitor colony growth, we image the plates using an array of document scanners, capturing colony expansion every 30 min for 5 days at 37^*°*^C (see Methods Sec. III for additional details). Our platform allows for direct control of local (resource competition) and nonlocal (initial founder arrangement) interaction strengths in characterizing intraand inter-colony structure in isogenic populations.

We extract colony boundaries using image analysis (see Methods Sec. III for details) and quantify singleand multi-colony structure as shown in Figure 1C. Anticipating that nutrient competition might play an important role, we also consider colony morphology, which is known to depend on nutrient availability [39, 40]. We compute two metrics: (1) inter-colony gap size used to determine whether colonies merge or form a stable gap and (2) colony roughness as a measure of morphology. Gap size is defined as the edge-to-edge distance between colonies, and roughness is quantified by the fraction of the convex hull not occupied by the colony, ℛ = 1 − *A*_colony_*/A*_hull_, so ℛ = 0 corresponds to a smooth colony front. These together serve as simple metrics that can be directly computed with any high-resolution imaging data.

### B. Intraand inter-colony structure is shaped by nutrient availability and founder deposition

How do these spatial structures depend on the initial arrangement of founder colonies and nutrient availability? Figure 2A–B shows the median final inter-colony gap size and colony roughness, respectively, measured across replicate colony pairs within bins of initial separation and supplied with the same initial BHI concentration (see Methods for binning details). Gap size exhibits systematic trends with both inputs, increasing with initial separation and decreasing with nutrient availability, indicating that this multi-colony metric depends on factors spanning cellular to colony scales (nonlocal). In contrast, colony roughness depends largely on nutrient level, with no clear dependence on inter-colony distance under the conditions surveyed here, indicating that this single-colony metric is governed primarily by cell-scale interactions (local). These results define a set of qualitative behaviors that any candidate model should reproduce.

**FIG. 2.**
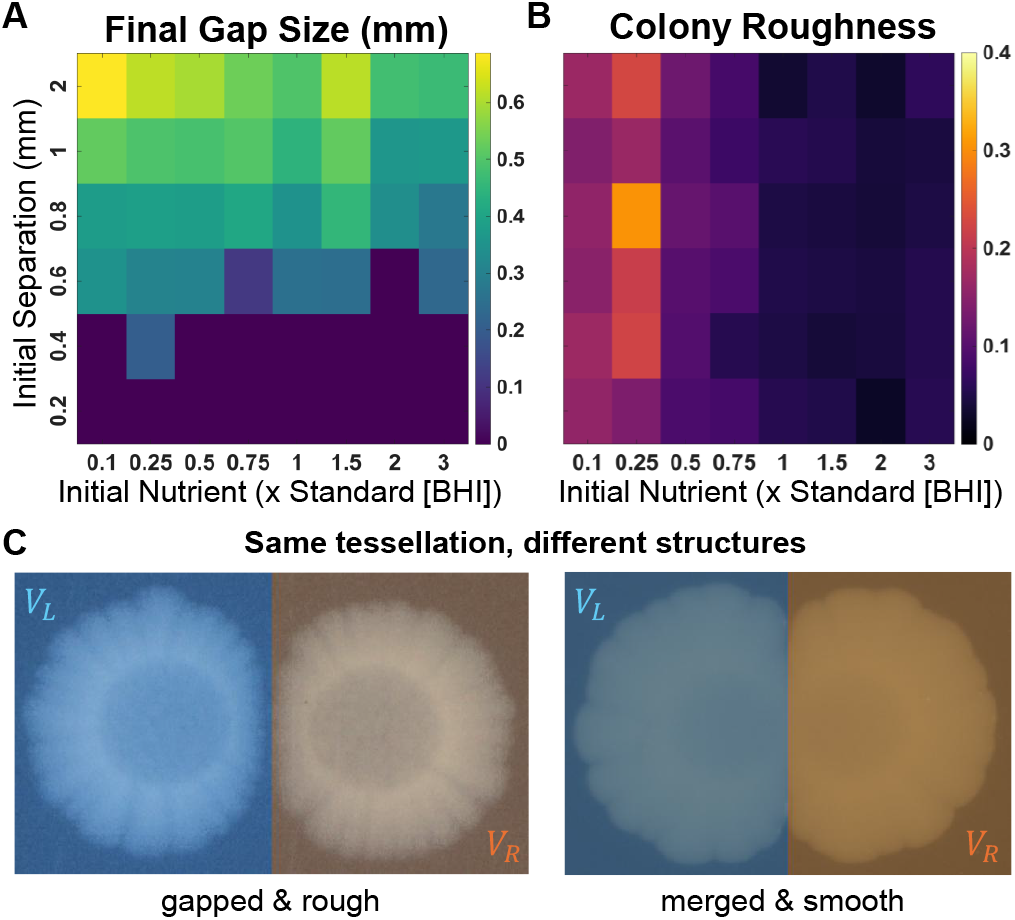
Spatial structures are determined by cross-scale interactions, not geometric constraints. **A:** Heatmap of final inter-colony gap size as a function of initial separation and nutrient level (fraction of standard BHI). Gap size increases with initial separation but decreases with nutrient availability. **B:** Heatmap of endpoint colony roughness from the same experiments as in (**A**). Roughness depends primarily on nutrient level, with no clear dependence on inter-colony distance. **C:** Representative colony pairs compared to a purely geometric Voronoi tessellation model (see text). In both cases, the Voronoi partitioning - and thus the spatial resource allocation - is identical, with left (*V*_*L*_) and right (*V*_*R*_) colonies assigned equal areas by the bisector set by their initial separation. Voronoi-based models therefore predict merging at the midline as each colony fills its domain. However, the experimental images show that such models fail to capture both colony morphology and the conditional merging (gap formation) observed between colonies. The left example corresponds to an initial separation of 1.5 mm and BHI fraction 0.1, and the right to 0.2 mm and BHI fraction 1. Colors in (**A**,**B**) denote median values within each bin aggregated over three independent experimental runs (see Methods).

As a baseline, we consider a purely geometric null model in which colony growth is limited only by the space available given the initial arrangement of founders. In this framework, space is effectively partitioned by a Voronoi tessellation, where each point in the domain is assigned to the colony to which it is initially closest, defining a Voronoi region for each colony. Under this assumption, colonies expand to fill their respective regions and meet at the boundaries set by the tessellation. Such models have been shown to capture coarse features of biomass partitioning across interacting colonies under certain conditions [30, 31]. However, Fig. 2C illustrates that identical tessellations can yield qualitatively distinct structures across both intra- and inter-colony scales. As a result, a Voronoi-based model is insufficient to describe the observed structures as it cannot distinguish between the presence or absence of inter-colony gaps, nor capture the emergence of colony roughness.

### C. Relating intra- and inter-colony phases to ecological interactions

Going beyond geometric constraints requires a dynamical description of growth. Experimental observations (Fig. 2A–B) indicate that nutrient competition plays an important role. Past work in other microbes suggests that inhibitory interactions are essential for such gap formation [28, 36]. But are these mechanisms necessary to capture our observed behaviors and how can we distinguish them? To answer this we build a hybrid agent-based model that generalizes the range expansion framework of Eden [41] by coupling population growth with reaction-diffusion dynamics of molecular fields. The model is defined on a square lattice with immotile cells occupying discrete sites at position (*x*_*i*_, *y*_*i*_) (Fig. 3A). Cells are assumed to proliferate by locally consuming a nutrient (*u*(*x*_*i*_, *y*_*i*_)), and secrete an inhibitor (*I*(*x*_*i*_, *y*_*i*_)) that in turn suppresses growth. Neglecting cell death, we consider a bacteriostatic inhibitor, which yields the birth rate

**FIG. 3.**
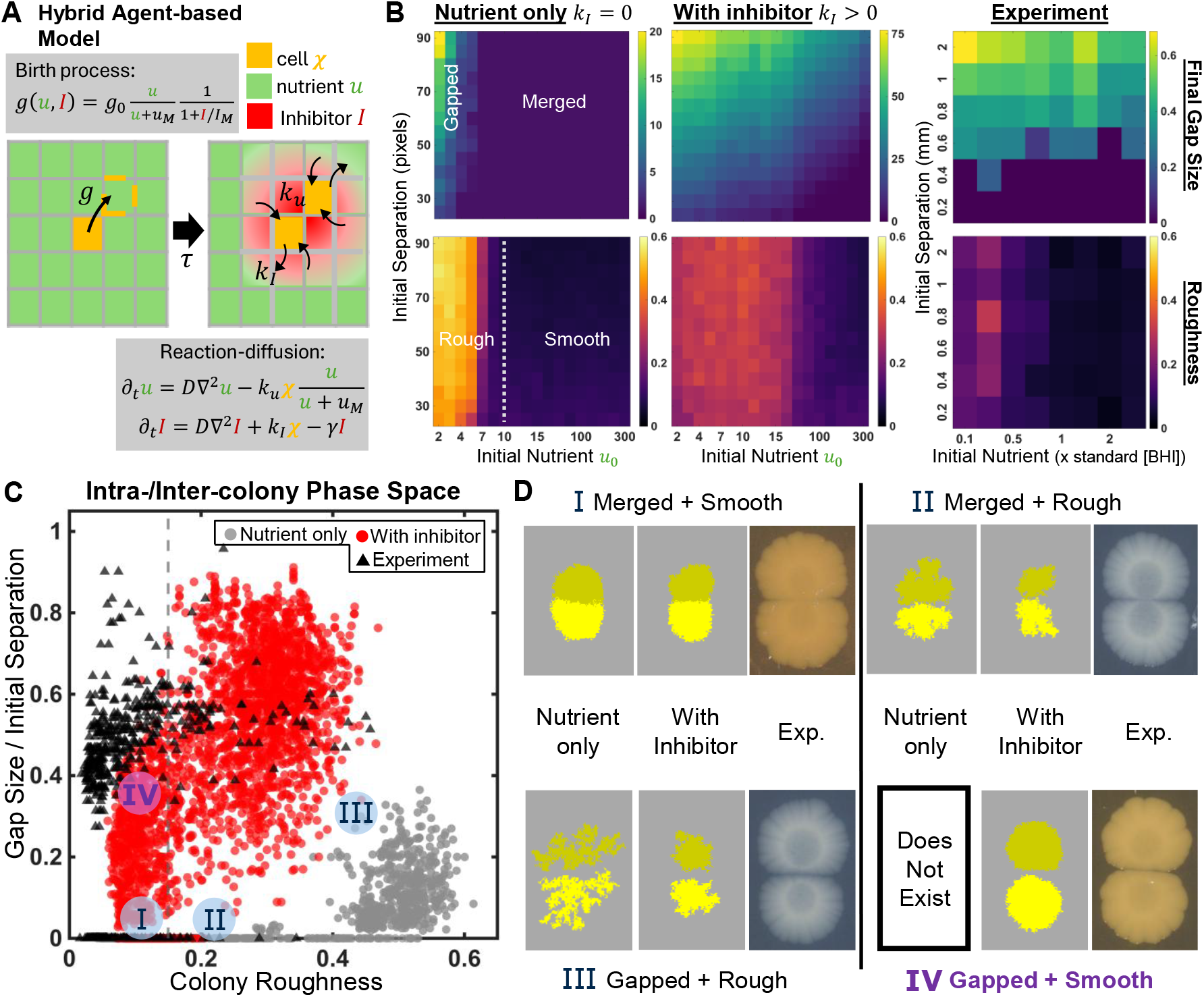
Model with resource competition and direct growth inhibition explains inter and intra-colony phases. **A:** Hybrid agent-based reaction–diffusion model of colony expansion. Cells on a lattice divide into neighboring sites with birth rates set by local nutrient *u* and inhibitor *I*, which are consumed (*k*_*u*_), produced (*k*_*I*_), and diffused (*D*) throughout space. Division requires mechanically pushing aside neighboring cells, mimicked by the screening factor *P* (see text) that prevents unphysical accelerating colony expansion (when *P* = 1, see SI). The model is initialized with a pair of cells seeded at varying initial separations and nutrient levels *u*_0_. We compare two scenarios: nutrient competition only (*k*_*I*_ = 0) and competition with direct growth inhibition (*k*_*I*_ = *k*_*u*_; see text).**B:** Heatmaps of final inter-colony gap size (top) and colony roughness (bottom) from simulations with nutrient-only interactions (left), with inhibition (middle), and experiments (right; replotted from Fig. 2). Each model pixel shows the median over 10 replicates per condition. The dashed line in the nutrient-only roughness panel indicates a known transition from smooth to rough morphology when *u*_0_ falls below a critical value set by consumption and division rates [39]. The model with inhibition qualitatively reproduces experimental trends in both metrics. **C:** Scatter plot of normalized gap size and colony roughness for individual replicates from (**B**). The dashed line delineates the morphological phases, defined from the smooth-to-rough transition of the nutrient-only model; see Supplemental Information (SI). The nutrient-only model (gray) occupies a distinct region from both experiments (black) and the model with inhibition (red), indicating that competition alone cannot reproduce the observed set of collective traits. Despite minor quantitative discrepancies due to finite size effects (see text and SI), the model with inhibition spans the full range of empirical intra- and inter-colony phases. **D:** Representative examples of each phase from simulations and experiments. The nutrient-only model captures three phases: merged and smooth, merged and rough, and gapped and rough, but fails to reproduce the fourth phase of gapped and smooth colonies observed in both experiments and the inhibition model. Simulated colonies are shown with two colors for the sake of visualization only.

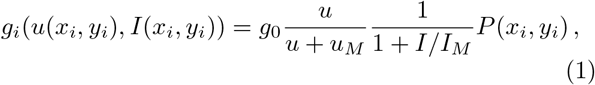

where *g*_0_ is the maximal growth rate and *u*_*M*_, *I*_*M*_ are Monod constants. *P* (*x*_*i*_, *y*_*i*_) ∈ [0, 1] is a screening factor that reduces the growth rate when the dividing cell would require pushing a long chain of neighboring cells. This factor is necessary to reproduce the constant radial expansion velocity commonly observed for growing colonies; without it (i.e., *P* = 1), we obtain exponential colony expansion (SI Fig. S2). We consider this as a phenomenological proxy for the friction-limited growth of mechanically compressible colonies [39, 42] (defined in Methods Sec. III E). Cell proliferation is simulated using a tau-leaping algorithm, where each cell undergoes a stochastic number of divisions with mean *g*_*i*_*τ* per time step *τ* ; division order and direction are weighted by growth rates and local occupancy (see Methods Sec. III E).

Simultaneously, the nutrient and inhibitor concentrations evolve according to the reaction–diffusion dynamics given by

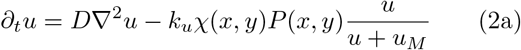

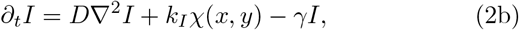

where *D* is the diffusion coefficient (assumed equal for both fields), *k*_*u*_ is the nutrient consumption rate, and *k*_*I*_ and *γ* are the inhibitor production and degradation rates, respectively. *χ*(*x, y*) indicates cell presence and *P* (*x, y*) is the same mechanical screening factor entering eq. 1 (so that nutrient consumption is suppressed identically to growth, conserving biomass in the absence of the inhibitor, while consumption is left uncoupled from growth when *I >* 0, consistent with known anabolism–catabolism uncoupling under growth inhibition [43]). These equations are numerically integrated at each time step to update the field values. Parameter values were chosen to match relative timescales without fitting to data (see Methods for details).

By varying a single parameter, the inhibitor production rate *k*_*I*_, we can explore regimes with (*k*_*I*_ *>* 0) and without (*k*_*I*_ = 0) any inhibitor. While the latter corresponds to nutrient competition alone (also recovered in the large *γ* limit; see SI Fig. S5), the former includes both competition and direct inhibition. For simplicity, we fix *k*_*I*_ = *k*_*u*_ when the inhibitor is included; deviations from this limit do not affect our results qualitatively. Upon simulating colony pairs across initial separations and nutrient levels, we obtain state diagrams (Fig. 3B) that show the transitions in the inter-colony gap size and colony roughness (quantified as in the experiments). By comparing these results for the two model variants with the experiments (replotted from Fig. 2), we see that both model variants reproduce most qualitative trends observed experimentally: gap size depends on both inputs, while roughness depends primarily on nutrient level. Consistent with prior work, the competition-only model exhibits a transition from smooth to rough morphologies below a critical nutrient level set by consumption dynamics [39] (dashed line in Fig. 3B). Together with the distinction between merged (gap size = 0) and gapped colonies, we identify four distinct collective “phases” that characterize intra- and inter-colony structure.

To compare phases directly, Fig. 3C plots all datasets in the plane of a normalized gap size (gap size divided by initial separation) versus colony roughness. In the competition-only model (gray), three phases emerge: (I) smooth, merged colonies at high nutrient levels; (II) rough, merged colonies at lower nutrient levels but small separations; and (III) rough, separated colonies with stable gaps at larger separations. Experimental data (black) populate all three phases (see Fig. 3D), but also reveal a fourth: (IV) smooth colonies that remain separated by a stable gap. This phase is inaccessible in the competition-only model but can be achieved when inhibitor production is included (red), yielding agreement across all four qualitative regimes. Fig. 3D shows examples of all phases of interacting colonies. We note that small quantitative discrepancies between model and experiment arise from finite system size effects in simulations, which limit the minimum achievable roughness; this is verified in SI Fig. S1.

These results establish that direct growth inhibition is necessary to reproduce the full phase space of experimentally observed behaviors. But how does a stable gap form in this case? Intuitively, gap formation reflects a race between two characteristic timescales: *τ*_front_ ∼Δ*/*2*v*, the time it takes both fronts to travel at speed *v* across half the initial separation Δ in the absence of any inhibitor, and *τ*_*I*_ *I*_*M*_ ∼*/k*_*I*_, the time for the inhibitor to locally accumulate till it arrests growth (neglecting degradation) leaving a gap of size *δ*. The fronts advance at a constant, mechanically-limited velocity which we compute to be *v* ≈ *g*_0_*L*_push_*κ*_*∞*_, where *L*_push_ is the width of the active growth layer (dependent on the screening factor *P*) and *κ*_*∞*_ a nutrient-dependent constant (see SI for details). The ratio of the two timescales defines a Damköhler-like number: Da = *τ*_front_*/τ*_*I*_ = *k*_*I*_ Δ*/*(2*g*_0_*I*_*M*_ *L*_push_*κ*_*∞*_) that captures the relative importance of front transport to a reaction rate. Fast moving fronts with Da ≪ 1 cause quick gap closure before the inhibitor can build-up. In contrast slow moving fronts with Da ≫ 1 allow sufficient accumulation of the inhibitor causing the fronts to arrest after traveling a distance of *vτ*_*I*_ and maintaining a stable gap of size *δ* ∼ Δ − 2*vτ*_*I*_ between colonies. To test this argument across a broad range of *k*_*I*_, *I*_*M*_, and inhibitor diffusivity, we simulate colony pairs using a one-dimensional reduction of the model (Fig. 4A), effectively focusing on the region near the gap midline between two large colonies, where the fronts are locally flat (see SI for details). Upon plotting the results in terms of Da and fitting a single constant Da_*c*_ ≈ 9.8 (the threshold for gap formation) in Fig. 4B, we find a striking data collapse of the final gap size onto the predicted curve, *δ* = Δ max(0, 1 −Da_*c*_*/*Da), in sharp contrast to the same data plotted against *k*_*I*_ alone (Fig. 4A). This result demonstrates that Da is the relevant non-dimensional parameter that controls gap formation of colonies. Direct tracking of the nutrient and inhibitor Monod factors at the gap midline in full two-dimensional simulations confirms that arrest is driven by local inhibitor accumulation rather than nutrient depletion at these operating points (SI Fig. S4).

**FIG. 4.**
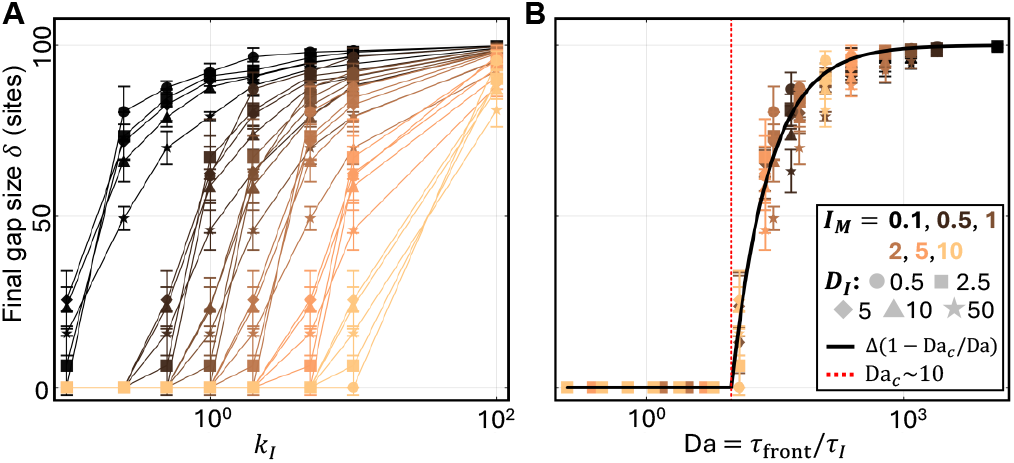
A ratio of timescales predicts colony merger and the final gap size. 1D simulations of colony pairs with varying inhibitor production rates *k*_*I*_, Monod constants *I*_*M*_, and diffusivities *D*_*I*_ (with *u*_0_ = 100, Δ = 100), while maintaining negligible degradation (*γτ*_front_ ≪ 1; see SI Fig. S5 for the general-*γ* case). **A:** Final gap size plotted against *k*_*I*_, for varying *I*_*M*_ (color), and *D*_*I*_ (marker shape). Onset of gap formation is delayed for larger *I*_*M*_ and is largely independent of *D*_*I*_ . **B:** Final gap size plotted against the Damköhler number Da = *τ*_front_*/τ*_*I*_ (see text) reveals data collapse onto the predicted curve : *δ* = Δ max(0, 1 − Da_*c*_*/*Da) (black curve). A single fit parameter Da_*c*_ ∼10 dictates the onset threshold (red dotted line) for the transition from colony merger (Da *<* Da_*c*_) to stable gap formation (Da *>* Da_*c*_). Symbols show mean ± standard deviation across 5 replicate seeds.

In summary, the ability to tune between competition-only and competition-plus-inhibition regimes allows us to assess whether resources alone can account for the observed phase behavior. The failure of the competition-only model to capture a key experimental phase indicates that an additional mechanism—direct growth inhibition—is required. We next test this prediction experimentally.

### D. Spent-agar test validates model prediction of gap formation from self-inhibition

To test whether *E. faecalis* colonies produce growth-inhibitory factors, we developed a simple spent-agar assay (Fig. 5A). Although we were unable to determine the molecular identity of the inhibitor, we aimed to establish the presence of growth inhibition at the ecological level, without resolving its specific mode of action. To establish this, we streaked a 2 cm ×2 cm perimeter from an overnight culture onto a standard BHI agar plate. After 5 days of growth, a ∼14 mm diameter disk was excised from the center of the spent agar within the square perimeter-only colony, where inhibitory factors are expected to accumulate (see Methods Sec. III C). The disk was placed in an empty well, supplemented with fresh nutrients (see Methods), and inoculated with fresh exponential-phase cells (∼10 CFUs*/μ*L; see Methods). Growth was then assessed after 24 hours. As a control, the same procedure was applied to initially blank agar plates lacking both nutrients and cells.

**FIG. 5.**
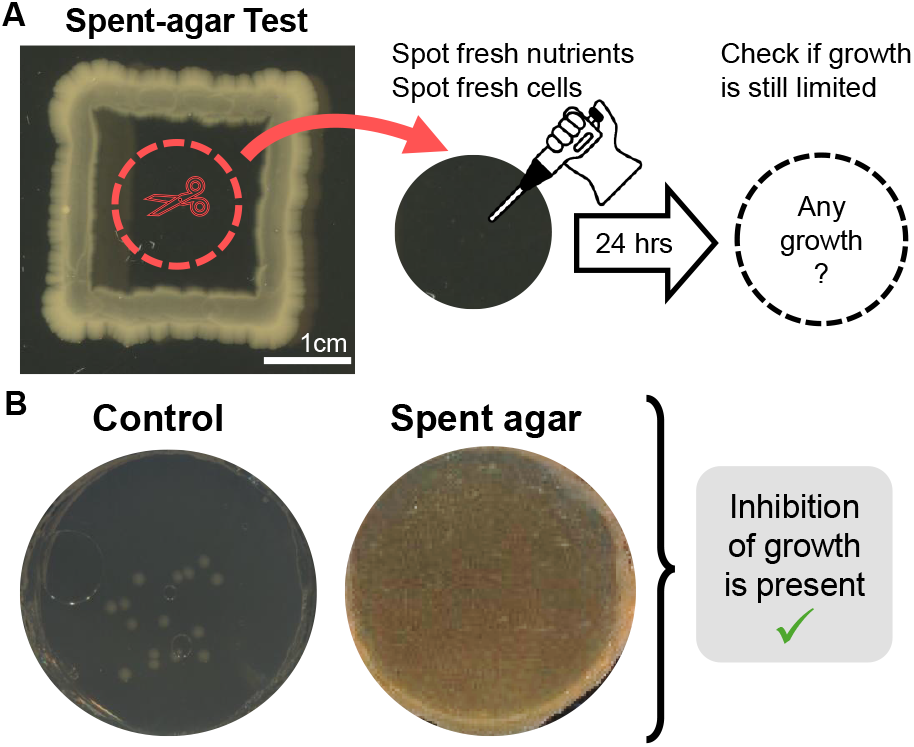
Spent-agar test validates model prediction of direct growth inhibition. **A:** Schematic of the spent-agar assay used to test the prediction that a direct growth inhibitor is present. Agar from the central region of square colonies grown for 5 days was excised and supplemented with fresh BHI (see Methods). The same procedure was applied to zero-nutrient control plates. Each disk was then inoculated with 2 *μ*L of fresh exponential-phase culture (∼ 15 cells) and imaged for 24 h. Because both conditions receive fresh nutrients, reduced growth on spent agar relative to control indicates the presence of a direct growth inhibitor. **B:** Representative endpoint images after 24 h for control (left) and spent-agar (right) conditions. The absence of visible growth on spent agar, compared to robust growth on control disks, confirms that direct growth inhibition is present alongside nutrient competition. See SI Fig. S6 for additional replicates.

This design isolates the effect of direct growth inhibition from nutrient depletion. Because both conditions receive fresh nutrients, reduced growth on spent agar relative to the control indicates the presence of inhibitory factors. Figure 5B shows representative endpoint images (see SI Fig. S6 for additional replicates): while robust growth is observed on control disks, no visible growth occurs on spent agar. These results demonstrate that direct growth inhibition is present in our system, consistent with the model prediction.

## II. DISCUSSION

By focusing on conditional gap formation in isogenic colonies (‘sibling rivalry’) as a minimal multi-colony trait, we have established a quantitative framework integrating high-throughput experiments, imaging, and modeling to investigate interacting bacterial communities with spatial structure. Using *E. faecalis* as a model system, we show how colony morphology and inter-colony gap formation reveal the presence of distinct ecological interactions mediated by diffusive processes. Our theoretical model captures the emergent single and multi-colony traits observed and identifies a mechanism beyond nutrient competition - specifically, direct growth inhibition - that must be present to explain all observations. An experimental test of this prediction using a spent agar assay validates the presence of an inhibitory agent whose accumulation drives gap formation between colonies.

Our robotic platform and *in-vitro* experiments allow for controlled, high-throughput, quantitative studies. But do similar behaviors occur in more natural environments? Several lines of evidence suggest that they do. Conditional biofilm merging patterns similar to those observed here can be seen in studies of colonization of the zebrafish gut [29], and inhibitory interactions are often more pronounced under stressful conditions [37], that are more likely in natural settings. Going beyond surface associated growth, roughness transitions analogous to those we observe have been reported in more complex, including three-dimensional, environments [44]. That sibling rivalry and antagonistic interactions are widely observed, including in non-laboratory strains e.g., soil isolates [28], suggests that the patterns and structural consequences we identify likely generalize beyond the controlled conditions studied here.

The minimal modeling framework captures the coarsegrained growth dynamics but neglects detailed physical (e.g., cell mechanics) and biological (e.g., cell death, phenotypic heterogeneity) effects that can be important. Future work incorporating these effects along with multiscale feedbacks will be needed to identify the detailed mechanisms underlying pattern formation [5]. Our spent-agar assay provides a simple and scalable diagnostic for identifying the types of interactions that govern spatial structure across these scales. But it only demonstrates the existence of an inhibitor without resolving the molecular agent or mechanism of action. That *E. faecalis* is known to produce lytic compounds (e.g., extracellular proteases) during biofilm formation [45] suggests potential inhibitory targets to be characterized in the future. The simplicity of our framework is nonetheless a strength as it identifies key collective processes essential for driving patterns, agnostic to specific molecular implementations, suggesting general principles that broadly apply across diverse systems.

In spatial settings, antagonistically interacting populations exhibit biological nucleation phenomena that are sensitive to initial conditions [38]. Our results suggest that self-inhibition behaves similarly, wherein the initial spatial configuration plays a central role in determining collective structure. A potential consequence of this dependence is that biofilm cohesion becomes strongly influenced by the spatial deposition of founder cells, with sparse inoculations leading to fragmented architectures. As biofilms exhibit greater antibiotic resistance compared to planktonic populations [46], this sensitivity to initial conditions points to a potential strategy for disrupting biofilm integrity by targeting early-stage spatial organization. Altogether, our quantitative study of sibling rivalry suggests how simple biophysical mechanisms can mediate non-specific recognition in bacteria [47], with implications for the evolution and ecological consequences of such interactions.

## III. METHODS

### A. Bacterial strains, media and culture conditions

All experiments were performed using *Enterococcus faecalis* strain OG1RF, a fully sequenced oral isolate [48]. Starter cultures were prepared by inoculating 5 mL of sterile REMEL™ Brain–Heart Infusion (BHI) broth (37 g L^*−*1^ in deionized (DI) water) from frozen stock and grown overnight at 37^*°*^C.

For multi-colony experiments, agar (1.5% [wt/vol] Fisher BioReagents™ Agar Powder/Flakes in DI water) supplemented with varying BHI concentrations was poured into 6-well plates (Corning™ Costar™), with 5 mL added per well. Due to the broad salt tolerance of *E. faecalis* [49–51], plates with ≤50% standard BHI were supplemented with 0.8% [wt/vol] NaCl to maintain osmotic balance and viability.

For spent-agar experiments, standard 100 mm petri dishes (Fisherbrand™) were prepared with either BHI agar (1.5% agar, 3.7% BHI [wt/vol]) for square-colony streak plates or agar (1.5% [wt/vol]) supplemented with 0.8% [wt/vol] NaCl for initially nutrient-free controls.

### B. Multi-colony growth experiment

Prior to inoculation onto agar plates with varying initial nutrient (BHI) levels (see Methods III A), inoculum properties were standardized across batches. Overnight cultures were diluted into fresh media to return cells to exponential phase, and inocula were prepared at OD_600_ ∼0.001.

Colony pairs were spotted using an Opentrons™ OT-2 liquid-handling robot, which dispensed 1 *μ*L droplets onto the agar surface at controlled initial separations. At small separations, droplets can merge prior to absorption, resulting in liquid-phase coalescence rather than colony-driven merging. To minimize such events, plates were pre-warmed at 37^*°*^C for ∼30 minutes to promote rapid absorption, and the robot protocol was designed to spot one colony per well across all plates before returning to deposit the second colony of each pair. Wells in which droplets merged or overlapped (e.g., due to resolution limits at small separations or variability in droplet size) were manually flagged prior to image processing and excluded from analysis.

Following inoculation, plates were sealed with black plate seals (Revity TopSeal-A™ Black) for contrast, parafilmed, and placed on flatbed document scanners (Epson™ Perfection V370) housed in a 37^*°*^C incubator. Colony expansion was imaged at a resolution of 1200 dpi every 30 minutes for 5 days. The combination of plate seals and parafilm provided sufficient moisture retention to prevent agar desiccation. Image acquisition was automated using the open-source software CMDTWAIN (www.gssezisoft.com).

Each scanner accommodates four 6-well plates, yielding 24 wells per scanner. With six scanners, a total of 144 wells were imaged per experimental run, each containing a pair of interacting colonies under specified nutrient and separation conditions. To ensure coverage of all conditions, three independent runs were performed, with each condition represented once per run.

### C. Spent-agar experiment

This experiment consisted of two groups: an experimental (spent-agar) condition and a control. For the experimental condition, cells from an overnight culture were streaked onto standard BHI agar plates in a 2cm ×2cm square perimeter and incubated for 5 days at 37^*°*^C after parafilming. Control plates, prepared with zero BHI as described in Methods III A, underwent the same incubation but without cell inoculation.

Following incubation, agar disks were excised from the central region of each square colony using a sterile 5 mL pipette tip (inverted to cut and transfer the disk) and placed into empty wells of a 6-well plate. The same procedure was applied to control plates to obtain non-spent agar disks. Each disk was then supplemented with two 15*μ*L droplets of concentrated BHI (37% [wt/vol]), applied sequentially to allow absorption. This ensured sufficient nutrient availability for subsequent growth without substantially diluting any accumulated inhibitory factors.

A 2 *μ*L inoculum containing ∼10 exponential-phase cells (prepared by diluting an overnight culture into fresh media) was then deposited onto each disk. Plates were sealed, parafilmed, and imaged as described in Methods III B, though only endpoint images were analyzed in this study.

### D. Image processing and analysis

Raw images acquired from flatbed document scanners were processed using custom Matlab scripts for preprocessing and segmentation. In total, 432 wells were monitored; wells were first identified and cropped from the full field-of-view, and those that were empty, contaminated, or contained overlapping colonies were excluded. After additional screening to remove wells affected by global agar surface shifts, 285 usable pairs of interacting colonies remained for analysis. We also note that brief scanner dropouts occurred during two runs but did not affect the reported results, which rely on early or late time points as described more below.

Processing of well images and extraction of intra- and inter-colony features were performed in Matlab. We developed a custom automated workflow that consisted of two pipelines – tracking inter-colony gap sizes and measuring colony morphology – each described more below. Both pipelines began with the same pre-processing steps to remove noise using standard techniques and built-in functions from Matlab’s image processing toolbox. The automated pipelines were able to generally capture the range of image properties (e.g., brightness, contrast, etc.) exhibited across well conditions and across time. In three cases wells required manual processing of the gap region.

For gap tracking and analysis, Otsu’s method was applied to denoised (cleaned and smoothed) images to segment colonies from the background [52]. The initial gap region was identified by computing the center-of-mass of each colony pair at early times, and analysis was restricted to zoomed-in regions around this gap for the full time course. Colonies were re-segmented in each zoomed frame to measure the initial gap size. Merging at the final time point was then assessed, defined as either (i) a single labeled object or (ii) a connected path of colony pixels spanning top to bottom. Trajectories identified as merged at the final time point were not further analyzed to reduce computational cost. Otherwise, the inter-colony gap size was quantified over time as the average thickness of the background region between colony masks; gap closure events were assigned a gap size of 0 and classified as merged.

For roughness measurements, homelands were identified at the initial time point and masked out from final segmentation masks, and colony pairs were split into top and bottom colonies for separate morphological analysis. Prior to Otsu segmentation, mean filtering was applied for contrast enhancement. Processing was performed in batch, with filtering parameters adjusted across conditions to account for contrast variability; accordingly, some segmentations required hole filling while others did not. Convex hull masks were generated from segmented colonies by filling boundary points, and roughness was computed as described in Results Sec.I A, with colony and hull areas given by the total pixels in their respective masks.

For visualization in the Results figures, the final gap size for each trajectory was defined as the minimum gap size attained over time. Gap size and roughness heatmaps were constructed by binning data according to the nominal BHI (nutrient) concentration of the plates and the measured initial separation, using bins of width 10 pixels for all but the largest separations, which were grouped into a final bin of width 40 pixels. With a scanner resolution of 1200 dpi used throughout this work, each pixel is approximately 21*μ*m.

### E. Resource competition and growth inhibition model

To simulate multi-colony expansion and interactions, we developed a hybrid agent-based model that generalizes the Eden growth model [41] by incorporating reaction–diffusion dynamics of nutrient and inhibitor fields, which modulate cell birth rates. We emphasize that our goal is not to reproduce fine-scale quantitative features, but to construct a minimal model that captures key qualitative observations: (1) inter-colony gap size depends on both founder arrangement and nutrient availability, whereas colony morphology depends primarily on nutrient level; and (2) inter- and intra-colony structure organizes into four phases – merged-and-smooth, merged- and-rough, gapped-and-rough, and gapped-and-smooth.

Simulations are initialized with two cells separated by a distance Δ on a square lattice of size *L* × *L* (with *L* = 250). The lattice state is represented by a matrix *χ*, where each site is either empty (0) or occupied (1). Nutrient *u* and inhibitor *I* are represented by corresponding *L* × *L* fields, initialized uniformly with *u*_0_ and 0, respectively. Each cell has a birth rate *g*_*i*_ (eq. 1) that depends on local *u* and *I*, and on the screening factor *P* (*x, y*) = Σ_*d*_ *w*_*d*_ exp(−*m*_*d*_*/L*_push_), summed over the lattice directions *d* accessible from (*x, y*): *m*_*d*_ is the number of contiguous occupied sites that would need to be displaced in direction *d* to reach an empty site, and *w*_*d*_ ∝ 1*/*(1+*m*_*d*_) (normalized over directions) biases divisions toward less-crowded directions. The same *P* (*x, y*) multiplies the nutrient consumption term (eq. 2a), so that a mechanically screened cell’s nutrient uptake is suppressed identically to its growth, conserving biomass; the limit *L*_push_ → ∞ recovers unconditional, infinite-range pushing. We adopt this screening as a phenomenological proxy to the friction-limited, mechanically compressible growth described in Ref. [39], where colony expansion is set by a competition between growth and mechanical relaxation rather than unconstrained pushing, since it is necessary to reproduce the constant-velocity colony expansion that unconstrained pushing fails to capture (SI Fig. S2).

Colony expansion is simulated using a *τ* -leaping scheme coupled to reaction–diffusion updates of *u* and *I*. The time step *τ* is set by the fastest process in the system, taken here as nutrient consumption, such that *τ* = 1*/k*_*u*_. Each iteration proceeds as follows:

1. For each cell, draw the number of division events from Poisson(*λ*_*i*_) with *λ*_*i*_ = *g*_*i*_*τ* .
2. Update nutrient and inhibitor fields by numerically integrating eq. 2 using an implicit diffusion solver with no-flux boundary conditions.
3. Apply division events sequentially in random order, weighted by *g*_*i*_:
  a. Each division targets a neighboring site chosen with weight 1*/*(1 + contiguous cell count), biasing against densely packed directions.
  b. If the target site is occupied, the chain of *m* contiguous cells along that ray is displaced outward by one lattice spacing with probability exp(−*m/L*_push_) (the same screening entering eq. 1); otherwise the division does not occur.
4. Repeat until a stopping condition is met:
  a. A colony reaches the lattice boundary, or
  b. Colonies merge or the inter-colony gap size remains unchanged for 10^3^ *τ* steps (gap size defined as the minimum boundary-to-boundary distance).

After simulation, colony roughness and final gap size are computed in the same manner as the experimental data and used in Fig. 3.

Parameter values were not fit to data but chosen to preserve relative timescales. We fix *g*_0_ = 1 and *u*_*M*_ = *I*_*M*_ = 1, and select remaining parameters to satisfy *g*_0_ *< D/l*^2^ *< k*_*u*_, *k*_*I*_, where *l* = 1 is the lattice spacing. Specifically, we set *D* = 5, *k*_*u*_ = 10, and *L*_push_ = 5, with *k*_*I*_ either 0 or *k*_*I*_ = *k*_*u*_ as described in Results Sec. I C. The inhibitor degradation rate *γ* is varied to probe three regimes: *γ* ∼*D/*(*L/*2)^2^ (long-range diffusion), *γ* ∼ *D/*(*L/*4)^2^ (inter-colony scale), and *γ* ∼ *D/*(*L/*100)^2^ (localized inhibition). The results in Fig. 3 use *γ* ∼ *D/*(*L/*2)^2^, while the other regimes are shown in the SI.

All simulations were performed in Julia using the Great Lakes High Performance Computing (HPC) cluster. The associated module code is available at https://github.com/jacobtmoran/MultiColonySimulator_Hybrid_ABM, and the full imaging, simulation, and assay data underlying this work are available at https://doi.org/10.7302/1fms-9074 (University of Michigan Deep Blue Data).

## Acknowledgments

We thank S. Kaplan, D. Lubensky, members of the Lubensky and Shankar groups, and the Wood and Zaman labs for helpful discussions. J.M. acknowledges the center for Advanced Research Computing (ARC) at the University of Michigan for providing computational resources through the Great Lakes HPC cluster to enable efficient simulation of models within this work. An AI language model (ChatGPT 5.1) was used to help edit phrasing of the text; all scientific content was written, verified, edited and approved by the authors. This work was supported in part by NIH Awards R35GM124875 (K.B.W.), RM1DE034220 (L.Z.), and R35GM166481 (S.S.). J.M. acknowledges funding support in part from the Michigan Pioneer Fellows Program. The content is solely the responsibility of the authors and does not necessarily represent the official views of NIH. S.S. also acknowledges partial support and helpful discussions at KITP (NSF PHY-2309135) and the Aspen Center for Physics (NSF PHY-2210452).

During the initial stages of this work, Kevin Wood unexpectedly passed away. Kevin was not only a passionate and creative scientist but also a great friend to many. J.M. is deeply grateful to Kevin for his inspirational mentorship during the genesis of this work.

## Author contributions

J.M. and K.B.W. conceived R. Porter, C. Trenado-Yuste, A. Martinez-Calvo, M. Su, and designed the multi-colony experiment. J.M. designed the overall research project, developed the empirical platform and modeling framework, performed all experiments, executed all simulations, processed and analyzed all data, and wrote the manuscript. M.S. was involved in preliminary experiments and discussions. M.H., S.S., R.J.W., and L.Z. provided guidance on all aspects of the research and edited the paper.

## SUPPLEMENTARY INFORMATION

## Appendix A: System size effects in morphology and delineating smooth versus rough

Overall, the model qualitatively recapitulates all empirical phases when inhibitor production is included, with remaining quantitative discrepancies arising from a lower bound on the achievable roughness (i.e., the vertical cutoff along the roughness axis beyond which model points do not extend in Fig. 3C). Here, we examine this limitation by showing that (1) the minimum roughness depends on colony (system) size, controlled by varying the simulation time *t* at a fixed lattice size (*L* = 250, matching the lattice size used throughout the main text), and (2) the smooth-to-rough transition is system-size independent, enabling consistent comparison between model and experiment.

To this end, we simulate single-colony growth using the competition-only model (where the smooth-to-rough transition is well characterized [39]) at fixed lattice size *L* = 250 and initial nutrient *u*_0_, with all other parameters as in Methods, recording colony morphology at several fixed simulation times *t* = 22, 26, 30, 34. Figure S1 shows the mean roughness as a function of *u*_0_ for each fixed *t*, demonstrating that the minimum achievable roughness decreases with increasing colony size (i.e., with increasing *t*). This behavior arises because larger colonies allow the smooth regime to approach *A*_colony_*/A*_hull_ →1 more closely, with fluctuations that diminish as colonies grow larger. In principle, allowing colonies to grow larger still (via larger *L* and/or longer *t*) would enable quantitative agreement with experiments; however, multi-colony simulations at sufficiently large system sizes are computationally prohibitive.

Importantly, the smooth-to-rough transition point is insensitive to system size, consistent with Ref. [39]. We therefore define the phase boundary using the stable inflection point of the roughness–*u*_0_ curves (ℛ ∼ 0.15). This value is consistent with the median roughness observed in simulations at *u*_0_ = *k*_*u*_*/g*_0_ = 10 and large initial separations, where colonies expand sufficiently before reaching the merge-based stopping criterion.

**FIG. S1.**
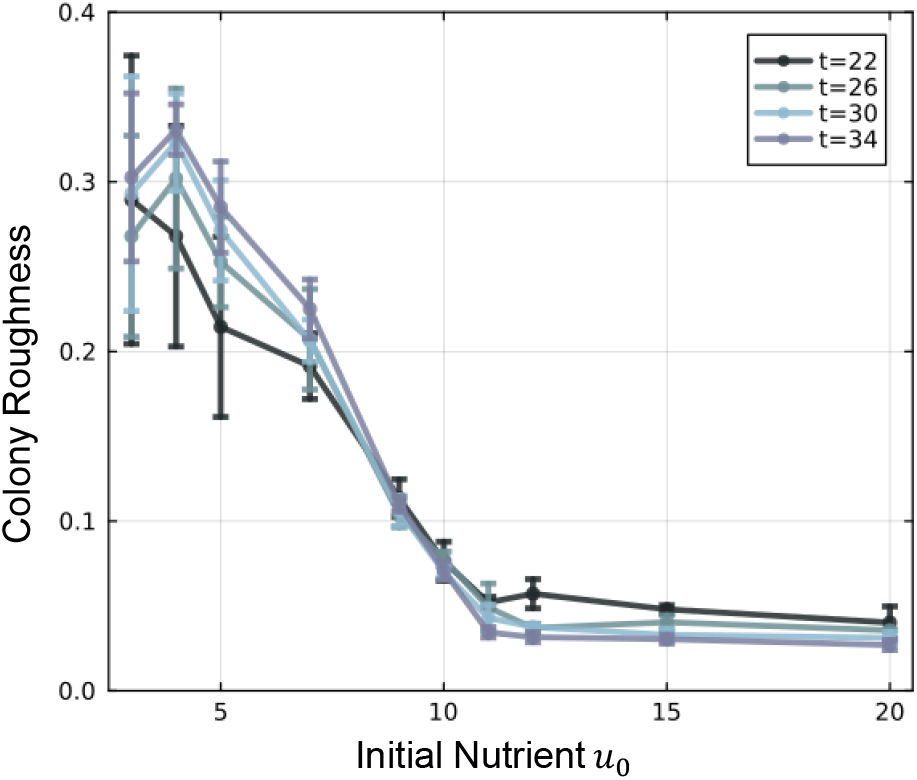
Minimum achievable roughness, but not the morphological transition, depends on colony size. Colony roughness (ℛ = 1 − *A*_colony_*/A*_hull_; see main text) is plotted as a function of initial nutrient *u*_0_ for single-colony simulations in the competition-only model, at fixed lattice size *L* = 250 (matching the lattice size used throughout the main text). Each curve corresponds to a different fixed simulation time *t*, which sets the achievable colony size. Markers and error bars denote the mean and standard deviation across 3 replicate seeds.

## Appendix B: Deriving and testing the timescale-race argument for gap formation

Here we derive the closed-form expression for *τ*_front_ used in the main text (Fig. 4) and the gap-size law it enters, and present additional numerical and trajectory-level evidence supporting the collapse.

The analysis in this section uses a one-dimensional reduction of the model: nutrient and inhibitor fields, and the two colony fronts, are represented along the single axis connecting the two colonies, rather than on the full two-dimensional lattice used elsewhere in this work (Methods Sec. III E). This is a natural simplification near the gap midline of two sufficiently large colonies, where the approaching fronts are locally flat and the relevant dynamics – nutrient depletion and inhibitor accumulation just ahead of each front – are effectively one-dimensional; curvature of the colony boundaries away from the midline does not enter. The reduction lets us efficiently sweep a much wider range of *k*_*I*_, *I*_*M*_, and *D*_*I*_ than would be practical in full two-dimensional simulations. The trajectory-level evidence for inhibitor-driven arrest presented below (Fig. S4) instead uses a full two-dimensional simulation, so that the same qualitative mechanism is confirmed independently of the one-dimensional reduction used for the quantitative collapse above.

Absent inhibitor, two colonies approach each other at a constant front velocity set by the mechanically screened growth rule introduced in the main text (Methods Sec. III E). Naive, unconditional pushing would instead predict an accelerating front, inconsistent with the friction-limited growth of mechanically compressible colonies [39]. We verify this directly for single-colony growth in Fig. S2: without screening (*P* ≡ 1), colony radius grows at an accelerated pace rather than at the constant velocity observed in the screened model (cell division succeeds with probability *P* exp(−*m/L*_push_)), confirming that the phenomenological proxy *P* ∼ (*x, y*) is necessary and sufficient for capturing this common empirical observation[10, 42, 53].

**FIG. S2.**
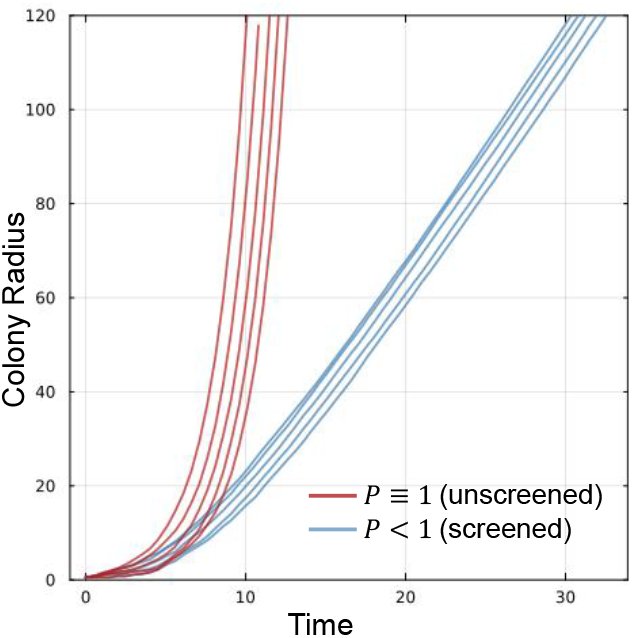
The screening factor *P* recovers constant-velocity colony expansion Colony radius for five individual replicate single-colony simulations (no inhibitor production, *k*_*I*_ = 0). Blue: screening as used throughout this work (*P* ∼exp(−*m/L*_push_), with *L*_push_ = 5), giving essentially linear radial expansion. Red: unconditional “pushing” upon division (*P* ≡ 1), which instead produces sharply accelerating growth. For the simulations, initial nutrient was set to *u*_0_ = 100 and colonies were seeded at the center of a square lattice of length *L* = 250.

The constant colony front expansion moves at a velocity *v* ≈ *g*_0_*L*_push_*κ*, where *κ* is the nutrient Monod factor *u/*(*u* + *u*_*M*_) averaged over the active growth layer. Per occupied site, division and nutrient-consumption rates are *g*_0_*f*_*u*_(*u*)*P* (*x, y*) and *k*_*u*_*f*_*u*_(*u*)*P* (*x, y*) respectively – the same mechanical screening factor *P* (*x, y*) entering eqs. 1 and 2a. Writing a steady, constant-velocity front and integrating the nutrient reaction-diffusion equation across the moving frame, the diffusive term vanishes (the fields are flat far ahead of and behind the front), leaving a flux balance *v*(*u*_0_ − *u*_*∞*_) = *k*_*u*_ *f*_*u*_(*u*) *P* (*ξ*) *dξ*, where *u*_*∞*_ is the nutrient concentration left behind the front. Because the front also advances due to the same divisions, *v* = *g*_0_ *f*_*u*_(*u*) *P* (*ξ*) *dξ* as well (so *L*_push_*κ* ≈ *f*_*u*_(*u*) *P* (*ξ*) *dξ*, with *L*_push_ the width over which *P* (*ξ*) is non-negligible and *κ* the corresponding *P* -weighted average of *f*_*u*_, recovering the earlier approximate *v* ≈*g*_0_*L*_push_*κ*), and the two expressions combine, exactly and independent of *D, L*_push_, *u*_*M*_, or the detailed shape of the active layer, to give

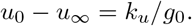

This also recovers the known nutrient-only stall (a front requires *u*_*∞*_ ≥ 0, i.e. *u*_0_ ≥ *k*_*u*_*/g*_0_ [39]) and pins the lower edge of the active layer’s nutrient throttle exactly, *κ*_*∞*_ ≡ *f*_*u*_(*u*_0_−*k*_*u*_*/g*_0_). Using *κ*_*∞*_ in place of the layer-averaged *κ* – an approximation that is exact deep in the nutrient-safe regime (*u*_0_ ≫ *k*_*u*_*/g*_0_) and improves as this margin grows – gives the closed-form estimate

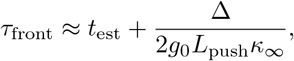

where *t*_est_ is a short colony-establishment offset and the factor of 2 accounts for both fronts closing the gap simultaneously. At the operating point used in Fig. 4 (*u*_0_ = 100), this closed-form estimate agrees with *τ*_front_ measured directly from nutrient-only simulations to within 2% (12.1 vs. 11.9 time units).

The one-parameter gap-size law used in the main text, *δ* = Δ(1 − Da_*c*_*/*Da), follows from the same timescale-race picture. Absent inhibitor, the gap remaining at time *t* is *δ*(*t*) ≈ Δ(1 − *t/τ*_front_). With inhibitor present, the local concentration in the still-closing gap builds up roughly as *k*_*I*_*t*, and growth is effectively arrested once this reaches the Monod scale *I*_*M*_, defining an arrest time *t*_arrest_ ∼ *I*_*M*_ */k*_*I*_ – the same combination entering the Damköhler-like number. Evaluating *δ*(*t*) at *t* = *t*_arrest_ gives *δ/*Δ ≈ 1 − 1*/*Da: the correct functional form, with a leading constant of exactly 1. The fitted constant Da_*c*_ absorbs everything this crude picture leaves out (e.g., front deceleration rather than constant velocity up to a sudden stop, inhibitor diffusion away from the gap, and production by both colonies into the same shrinking gap, etc.) without changing the functional form. Because the same expression predicts merging exactly when Da = Da_*c*_, we use the same fitted Da_*c*_ for both the merge threshold and the gap-size law in Fig. 4, rather than fitting them independently.

To test this argument, we simulated colony pairs at fixed *u*_0_ = 100 and Δ = 100 across a wide range of *k*_*I*_, *I*_*M*_, and inhibitor diffusivity *D*_*I*_, restricting to conditions where inhibitor degradation is negligible over the closing time (*γτ*_front_ *<* 0.1; the general-*γ* dependence is addressed separately in Fig. S5 at the level of qualitative phase structure). Figure S3 shows that this collapse is essentially identical whether *x* is built from the measured (Da_*c*_ ≈ 9.7) or the closed-form (Da_*c*_ ≈ 9.8) *τ*_front_ (*R*^2^ = 0.956 in both cases), confirming that the closed-form estimate used in the main text (Fig. 4) performs no worse than directly measuring *τ*_front_ from nutrient-only simulations. As already shown in the main text (Fig. 4A), this agreement is specific to the combined variable *x*: plotting the same data against *k*_*I*_ alone, without dividing by *I*_*M*_, eliminates the collapse entirely (*R*^2^ = −0.184), with each *I*_*M*_ tracing out its own separate trend.

**FIG. S3.**
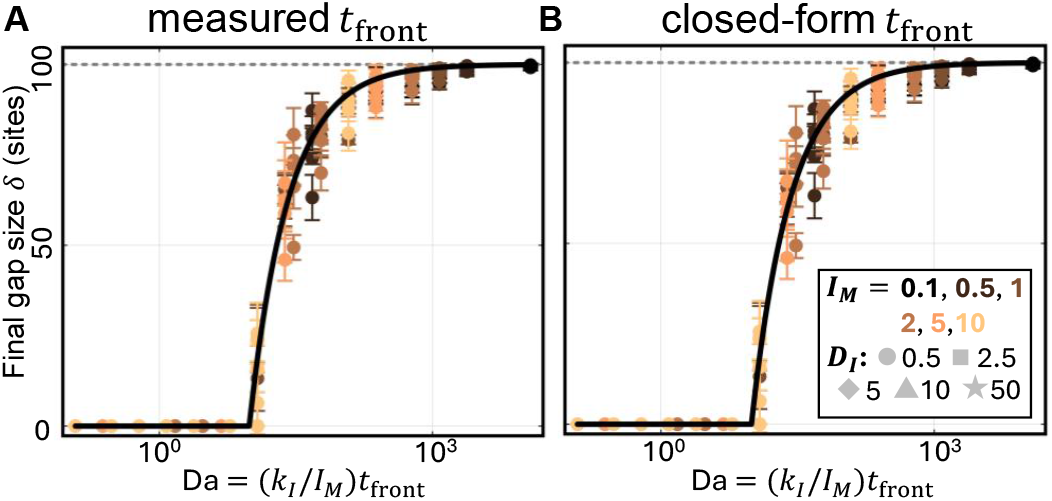
The closed-form *τ*_front_ is a good approximation compared against the exact *τ*_front_ measured directly. Final gap size vs. Da = (*k*_*I*_ */I*_*M*_) *τ*_front_, using the directly measured (**a**) or closed-form (**b**) *τ*_front_; both collapse equally well onto the fitted law *δ* = Δ max(0, 1 −Da_*c*_*/*Da) (black curve), with the same fitted threshold Da_*c*_ in both panels. Compare to the un-collapsed control plotted against *k*_*I*_ alone in Fig. 4A.

Finally, we obtained direct trajectory-level evidence for which mechanism is responsible for arrest by tracking the nutrient and inhibitor Monod factors, *f*_*u*_ = *u/*(*u* + *u*_*M*_) and *f*_*I*_ = *I*_*M*_ */*(*I*_*M*_ + *I*), at the gap midline over the course of a full two-dimensional simulation (Fig. S4). At *u*_0_ = 100 (the operating point used throughout this section), the front stabilizes with *f*_*u*_ essentially unchanged from its initial value (a 2.6% drop) while *f*_*I*_ has collapsed by 98%, directly confirming inhibitor-driven arrest rather than local nutrient depletion.

**FIG. S4.**
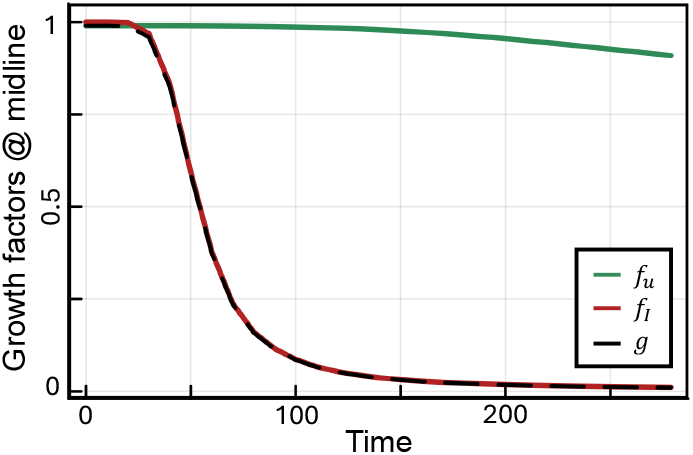
Direct trajectory evidence for inhibitor-driven arrest. Nutrient (*f*_*u*_, green) and inhibitor (*f*_*I*_, red) Monod growth factors, and their product *g* = *g*_0_*f*_*u*_*f*_*I*_ (dashed), tracked at the gap midline over time in a full two-dimensional simulation (*u*_0_ = 100, *k*_*I*_ = 10, *I*_*M*_ = 5). *f*_*I*_ collapses well before *f*_*u*_ moves appreciably, confirming that growth arrest here is driven by local inhibitor accumulation rather than nutrient depletion.

## Appendix C: Increasing rate of inhibitor degradation is consistent with nutrient-only limit

To assess the robustness of the model predictions to inhibitor dynamics, we varied the inhibitor degradation rate *γ* across regimes spanning long-range to highly localized inhibition (Fig. S5). Specifically, we consider values of *γ* corresponding to characteristic diffusion lengths that are system-wide, inter-colony scale, and confined to individual colonies (see Methods).

Across these regimes, the qualitative phase structure remains largely unchanged so long as inhibition acts over distances comparable to or larger than the inter-colony separation. In particular, the model continues to exhibit all four intra- and inter-colony phases identified in the main text, including the gapped-and-smooth regime that cannot be captured by nutrient competition alone. This demonstrates that the emergence of this additional phase is not sensitive to the precise details of inhibitor transport or lifetime, but rather to the presence of sufficiently long-ranged inhibitory interactions.

**FIG. S5.**
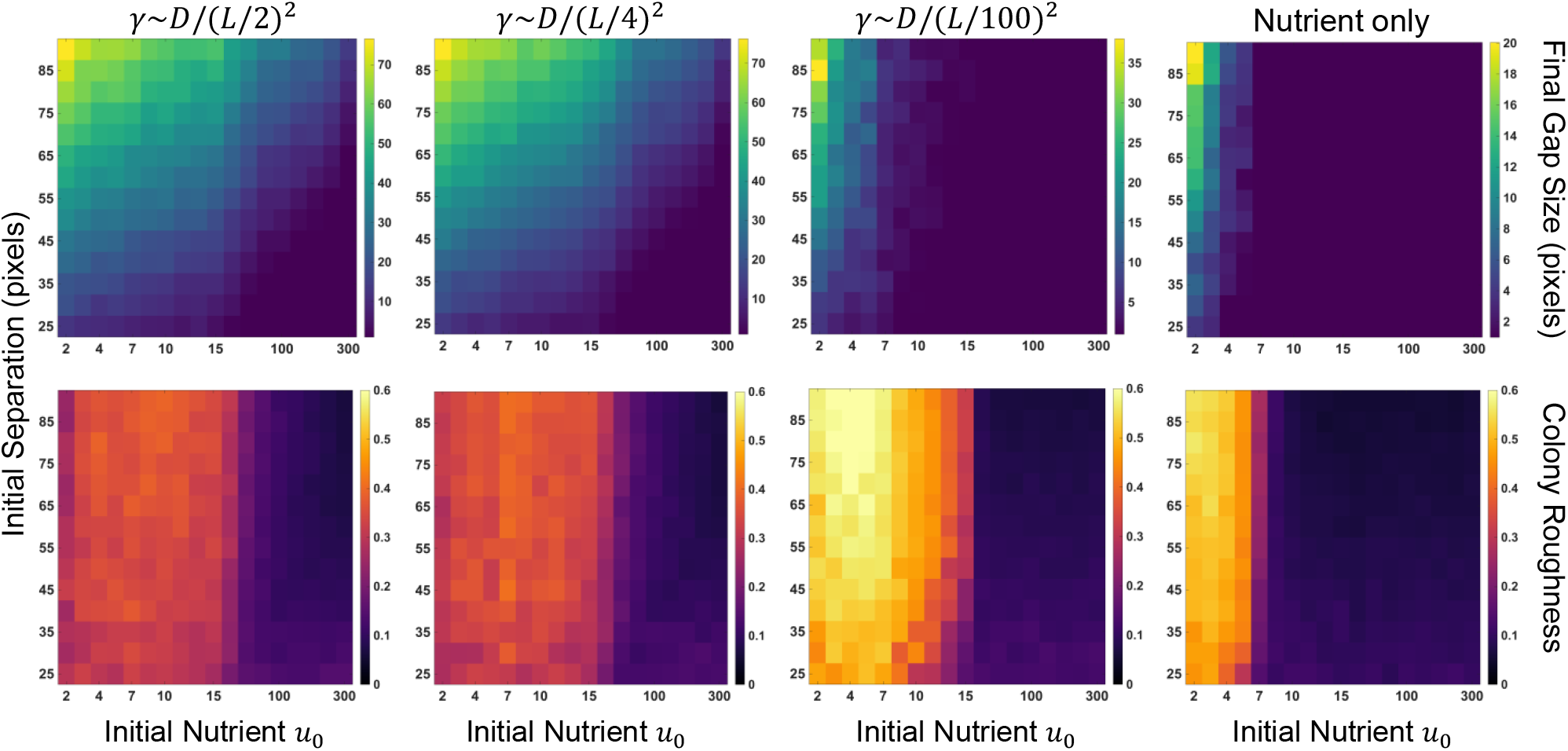
Localized inhibition recovers the nutrient-only limit. Heatmaps of final inter-colony gap size (top) and colony roughness (bottom) as functions of initial separation and nutrient level *u*_0_ from colony-pair simulations (see Methods Sec. III E). Columns correspond to increasing inhibitor degradation rate *γ*, with the rightmost column showing the nutrient-only model (*k*_*I*_ = 0). The degradation rate is expressed via the characteristic diffusion length scale *L*_*I*_ relative to the system size *L* = 250: *γ* ∼ *D/*(*L/*2)^2^ (*L*_*I*_ = 100, left) corresponds to system-wide inhibitor spread, *γ* ∼ *D/*(*L/*4)^2^ (*L*_*I*_ = 50) to an inter-colony scale, and *γ* ∼*D/*(*L/*100)^2^ (*L*_*I*_ = 3) to strongly localized inhibition. As degradation increases, the model recovers the three-phase structure of the nutrient-only limit, losing the additional phase of smooth colonies separated by persistent gaps. These regimes show that the qualitative phase structure is robust to inhibitor dynamics while recovering intuitive limiting behaviors. Each heatmap pixel reports the median across 10 replicate simulations.

In contrast, when degradation is sufficiently rapid such that inhibition becomes tightly localized to individual colonies, the model reduces to the nutrient-competition-only limit. In this regime, the phase space collapses to three phases, losing the gapped-and-smooth configuration and recovering the same qualitative behavior as the *k*_*I*_ = 0 case.

Together, these results show that the model predictions are robust across a range of biologically relevant inhibitor dynamics, while also recovering the expected limiting behavior when inhibition cannot mediate inter-colony interactions.

**FIG. S6.**
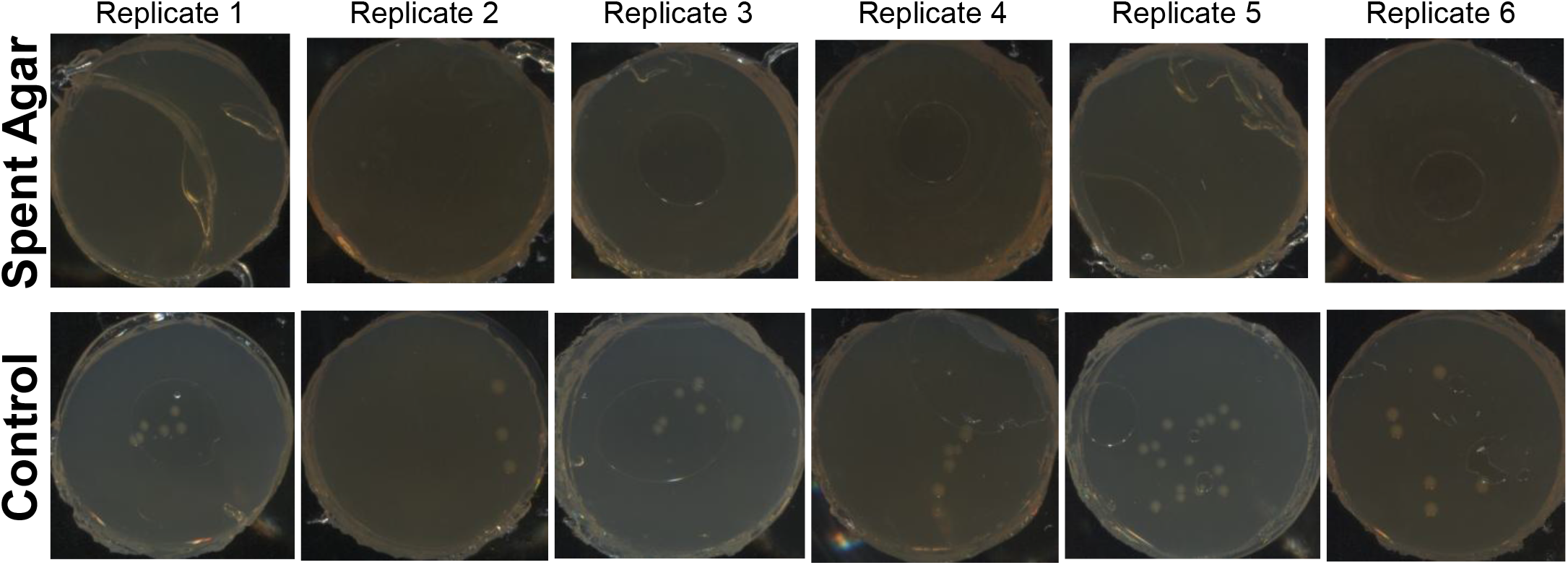
Additional replicates from spent-agar test. Top row shows images of replicate spent-agar disks excised from the center of a colony streaked in a square perimeter as described in the Methods. Similarly, the bottom row shows agar disks from the control (non-spent) condition. All images were acquired at the 24 hour time point of incubation. Comparing the two sets of replicates it is clear that there is no visible colony growth on the spent-agar disks relative to the control disks, indicating the presence of a direct growth inhibitor in the spent agar (see text).

